# Assembly of a Parts List of the Human Mitotic Cell Cycle Machinery

**DOI:** 10.1101/280339

**Authors:** Bruno Giotti, Sz-Hau Chen, Mark W. Barnett, Tim Regan, Tony Ly, Stefan Wiemann, David A. Hume, Tom C. Freeman

## Abstract

The set of proteins required for mitotic division remains poorly characterised. Here, an extensive series of correlation analyses of human and mouse transcriptomics data was performed to identify genes strongly and reproducibly associated with cells undergoing S/G2-M phases of the cell cycle. In so doing, a list of 701 cell cycle-associated genes was defined and shown that whilst many are only expressed during these phases, the expression of others is also driven by alternative promoters. Of this list, 496 genes have known cell cycle functions, whereas 205 were assigned as putative cell cycle genes, 53 of which are functionally uncharacterised. Among these, 27 were screened for subcellular localisation revealing many to be nuclear localised and at least four to be novel centrosomal proteins. Furthermore, 10 others inhibited cell proliferation upon siRNA knockdown. This study presents the first comprehensive list of human cell cycle proteins, identifying many new candidate proteins.

## Introduction

Mitotic cell division is a process common to all eukaryotic organisms and achieved through a highly orchestrated series of events classified into four sequential phases: G_1_ (gap phase), S (DNA replication), G_2_ and M (mitosis). The concerted action of hundreds of proteins is required to drive the process through to a successful conclusion, many of which are expressed in a phase-specific manner. They mediate processes such as DNA replication and repair, chromosome condensation, centrosome duplication and cytokinesis. Dysregulation or mutation of genes encoding proteins essential for high fidelity DNA replication, is often associated with disease, in particular cancer^1,2^. Accordingly, known components of this system are important therapeutic targets^3^ and novel components might present new therapeutic opportunities.

Many of the key proteins required for mitotic division are known from studies in model organisms including yeast, as well as mammalian cells^4,5^. With the aim of identifying all the components of the system, high content analysis platforms have been utilised. For example RNAi screens^6-8^, CRISPR/Cas9^9^, proteomics ^10,11,12^ studies have all proposed lists of cell cycle genes/proteins but a consensus between studies has not emerged. In particular, genome-wide transcriptomics studies^13-19^ have identified sets of transcripts sequentially regulated during the cell cycle phases in multiple species but comparison of results from four studies of different human cell lines identified only 96 genes in common^15^. Our reanalysis of these data, taking account of some of the technical variables, suggested that the true concordance of the cell cycle transcriptional network across cell types is much greater^20^. Furthermore, our analyses of large collections of tissue and cell transcriptomics data commonly identify a large cluster of cell cycle transcripts whose expression is elevated in cells or tissues with a high mitotic index^21-25^.

We report here on a data driven curation exercise aimed at identifying the cohort of genes up-regulated in all human cell types from the G_1_/S boundary through to the completion of M phase (S/G2-M). There are of course, many growth-associated transcriptional regulatory events including activation of cyclins, cyclin-dependent kinases and E2F transcription factors, that occur during G1 and are a precondition for entry into S phase^26^, but these are not the focus of this study. We monitored genome-wide gene expression in primary human dermal fibroblasts (NHDF) cells as they synchronously enter the cell cycle from a resting state (G0). Using network co-expression analysis and clustering of the data, we identified a cell cycle-enriched cluster associated with S/G2-M phases. To refine this initial list, we identified those genes that were robustly co-expressed when their transcription was examined across multiple different human primary cell types in the promoter-based FANTOM5 transcriptional data set^27^ and in synchronised murine fibroblasts. Manual curation of these data resulted in a list of 701 genes strongly associated with the S/G2-M phase transcriptional network. Of these, 496 encode proteins with known functions within cell division, 145 of which were not identified in any of the previous human cell cycle transcriptomics studies. Of the remaining 205 genes, 53 encode functionally uncharacterised proteins. To further validate this discovery set, we examined their expression across a range of human tissues and in mouse embryonic tissues during development. We also performed functional assays including protein localisation by over-expression of GFP-tagged proteins and knockdown by RNAi.

## Methods

### Cell culture and synchronization

Primary human dermal fibroblasts (NHDF) isolated from neonate foreskins (gifted by Dr Finn Grey, University of Edinburgh, UK) were plated on 175 cm^2^ tissue culture flasks (Thermo Fisher, Perth, UK) at density of 6×10^3^ cells/cm^2^. Cells were cultured in DMEM (Sigma-Aldrich, Missouri, US) with 10% (v/v) foetal calf serum (FCS) (GE Healthcare, Little Chalfont, UK) and antibiotics (25 U/ml penicillin and 25 μg/ml streptomycin (Life technologies, Paisley, UK). Starvation-induced synchronisation was achieved by replacing full medium with DMEM containing 0.5% FCS for 48 h in accordance with published methods^28^. After this time, medium was replaced with DMEM containing 10% FCS promoting the synchronised entry of the NHDF back into the cell cycle. Similarly, mouse embryonic fibroblasts (MEF) were cultured in 175 cm^2^ tissue culture flasks (Thermo Fisher) at a density of 6,000 cells/cm^2^ in DMEM with 10% FCS and the same protocol followed as for NHDF synchronisation.

In both cases, cell synchronisation was assessed after 48 h of serum starvation and at various time points following the re-addition of complete medium using a BD LSR Fortessa X-20 flow cytometer (BD Biosciences, San Jose, CA, US) with propidium iodide staining. Unsynchronised populations were evaluated to assess the degree of synchronisation achieved. For protein localisation assays, 1×10^5^ HEK293T cells were grown in DMEM (Sigma-Aldrich) medium plus 10% FCS, 1% Glutamax (Gibco, Gaithersburg, US), 1% non-essential amino acids (Gibco) and 25 Units/ml penicillin/streptomycin on 13 mm glass coverslips previously coated with poly-L-lysine (0.1 mg/ml) in each well of a 24-well plate. Cells were grown until coverage of approximately 70% was obtained. To increase percentage of cells undergoing mitosis, HEK293 were reversibly blocked at the G_2_/M boundary with RO3306, as described previously^29^.

### Microarray preparation

Two human microarray datasets were generated using normal human foreskin fibroblasts (NHDF). For the first microarray experiment duplicate samples were taken at 6 h intervals over a 48 h period (24 samples in total including unsynchronised control cells cultured in parallel and harvested at 0 and 24 h). In a second independent experiment, samples were collected at 1 and 2 h following re-addition of complete medium, and then every 2 h for a 24 h period (16 samples in total including two control samples). In a third microarray experiment using mouse fibroblasts, MEF samples were collected at 0, 0.5, 1, 2 h following re-addition of complete medium and then every 2 h for a 30 h period (24 samples in total including replicates for 0, 12, 18, 24 h samples and unsynchronised control samples). For all experiments described above total RNA was isolated using the RNeasy Mini Kit (QIAGEN, Manchester, UK) according to manufacturer’s instructions. cDNA was generated by the reverse transcription of total RNA (500 ng) using the Ambion WT Expression Kit (Life technologies), fragmented and then labelled by TdT DNA labelling reagent using GeneChip^®^ WT Terminal Labelling Kit (Affymetrix, Buckinghamshire, UK) according to manufacturer’s instructions. Samples were hybridized to Affymetrix Human Gene 1.1-ST Arrays for both NHDF time course experiments and to the Mouse Gene 2.0 ST Arrays for the MEF experiment using an Affymetrix GeneAtlas system. For cross-validation the Fantom5 (F5) primary cell atlas of human promoter expression^21^ was used, including 495 samples from about 100 human primary cell types. The data is publicly available at http://fantom.gsc.riken.jp/5/ ^30^.

### Data pre-processing

Raw data (.cel files) derived from the three microarray experiments were pre-processed using *Bioconductor* (www.bioconductor.org). The package *ArrayQualityMetrics* was used to perform QC on the data. All arrays passed the various tests carried out by the package and expression levels were normalised using Robust Multiarray Averaging (RMA) normalisation using the *Oligo* package. The two normalised NHDF datasets were also adjusted with the batch correction algorithm ComBat^31^ to adjust for variations in the average intensity between experiments. Low intensity signal probesets (< 20) were removed (a total of 9,408 probesets). Likewise, a filtering of low-end signal was applied on the FANTOM5 primary cell atlas removing promoters with <5 tags per million (TPM) reads. Probe set annotations were retrieved with the *hugene11transcriptcluser.db* package for the human data and with *mogene20sttranscriptcluster.db* package for the mouse data. Mouse to human orthologues were retrieved from the web resource Mouse Genome Informatics (MGI) (http://www.informatics.jax.org/).

### Network analysis

The NHDF, FANTOM5 (primary cell and mouse development datasets)^27^, Tissue atlas dataset^32^ and MEF datasets were subjected to network-based correlation analyses. Data was loaded into the tool Miru (Kajeka Ltd., Edinburgh, UK) and Pearson correlation matrices were calculated comparing expression profiles between individual samples or genes, and these were used as the basis to construct co-expression networks as described previously ^23^. Correlation thresholds for all analyses were set to allow minimal contribution of random correlations to the analyses. These were based on a comparison of the correlation distributions of the experimental datasets vs. permuted measurements from 2,000 randomly selected measurements. Values selected also minimised the number of edges whilst maintaining a maximum number of nodes (Figures S1-2). The two NHDF time-course experiments were analysed together. A correlation network was constructed using a threshold of *r* ≥ 0.88 and the graph clustered to identify co-expression modules of genes with a broadly similar expression pattern using the MCL clustering algorithm^33^ with the inflation value (which controls the granularity clustering) set to 1.3 (MCLi = 1.3). Clusters of genes whose expression varied for technical reasons, i.e. profile associated with a batch or experiment, were removed. The correlation network of the remaining data comprised of 4,735 nodes (probesets) connected by 153,809 edges. Using different inflation values the network was divided into a few (MCLi 1.3) or many (MCLi 2.2) clusters of transcripts. Transcripts within the S/G2-M cluster (cluster 3) plus all nodes immediately adjacent to them (n+1), were then selected for further analysis. The node walk expansion was to capture a number of similarly expressed genes on the periphery of the main cluster. Entrez IDs of the cell cycle-associated transcripts identified in the NHDF data were used to subset the FANTOM5 primary cell atlas, prior to network analysis. The subset FANTOM5 primary cell atlas data was then parsed at *r* ≥ 0.5 and clustered at MCLi = 1.7. The MEF time-course data was parsed at *r* ≥ 0.88 and the resultant networks clustered using MCLi = 2.2. The Tissue Atlas and FANTOM5 mouse development datasets were subset for the curated S/G2-M gene list, plotted at r ≥ 0.5, and clustered at MCLi 3.2 and 1.7, respectively.

### Assembly of evidence and annotation of the cell cycle ‘parts list’

In assembling a list of cell cycle genes we have sought to bring together various sources of evidence to support this association. This includes whether they were implicated by the current studies of their expression in experiments performed on NHDF, MEF or human primary cell atlas, previous transcriptomics studies on human cells^13,14,16,18,34,35^, the Mitocheck database^36^ and human protein atlas (HPA)^37^ resource, and finally, our own functional assays. Furthermore, we examined evidence from the literature as well as annotation and pathway resources to provide, where possible, a functional grouping and annotation for each gene. This was carried out by retrieving UniprotKB biological process terms (UniprotKB *keywords*) and when none were found for a given gene, annotation was supplemented from other sources, namely Gene Ontology^38^ and Reactome^39^. These efforts were backed up by extensive review of the published literature. The full list of genes with their corresponding functional annotation can be found in S1 Table. Based on this work, genes were further classified based on evidence of their involvement in cell cycle: The *Known* group defines genes for which there is robust evidence of their involvement in one of the pathways associated with the cell cycle, whereas the *Putative* group includes genes for which there is little or no direct evidence of them being involved in the cell cycle. This group also includes a number of functionally uncharacterised genes. Finally, a simple confidence score was used to order the cell cycle list based on the weight of evidence supporting a gene’s involvement in the cell cycle; One point was awarded to all genes for each line of evidence supporting their association with the cell cycle, i.e. they were identified by the current or five previous human transcriptomics studies^13,14,16,18,34^, their knockdown generated a mitosis-related phenotype in the current Mitocheck screen^36^ and whether the gene has been associated with a cell cycle-related phenotype in human and mouse^40,41^.

### Gene Ontology and motifs enrichment analysis

GO enrichment analyses were conducted with the Database for Annotation, Visualization and Integrated Discovery (DAVID, v6.8), a web-based tool for Gene Ontology enrichment analysis (http://david.abcc.ncifcrf.gov/). Gene sets within the clusters generated by the MCL algorithm were analysed for GO_BP terms using the Functional Annotation clustering tool. Motifs enrichment analysis was conducted using HOMER^42^ through the CAGEd-oPOSSUM web tool^43^. Genomic loci of the cell cycle-associated promoters were inputted in the software. Enrichments for known motifs were searched between 1,000 bp upstream and 300 bp downstream from the TSS.

### RTCA analysis following gene knockdown by RNAi

NHDF cells were cultured in Dulbecco Modified Eagle Medium) DMEM, Sigma-Aldrich (with 10) %v/v (foetal bovine serum (FBS, GE Healthcare) and 25 U/ml penicillin and 25 μg/ml streptomycin (Life technologies). The xCELLigence (Roche, Penzberg, Germany) real time cell analyser (RTCA) system was used to monitor the effect of gene knockdown on cell impedance, taken as a proxy for cell proliferation. Background impedance for the E-plates 96 (ACEA Biosciences, San Diego, US) was standardized by the addition of culture medium (DMEM with 10% FBS, and 25 U/ml penicillin and 25 μg/ml streptomycin) following the manufacturer’s instructions. Following trypsination, cells were seeded at density of 6,000 cells/cm^2^ in each well of the 96 well E-plates with the additional of 100 μl complete medium. Baseline levels of cell impedance index recorded and 24 h later, esiRNA transfection esiRNAs (Sigma-Aldrich) was performed while plates were undocked from the RTCA station. Transfection of esiRNA was carried out using the transfection reagent SilenceMag (OZ bioscience, Marseille, France). esiRNA was combined with 3.3 μl SilenceMag and 3.0 μl H_2_O, and then mixed with antibiotic-free medium in a final volume of 100 μl and a concentration of 50 nM esiRNA per well. Complete medium was then replaced with the transfection mix, placed on magnetic plates (OZ bioscience) for 30 min in the incubator under the condition of 5% CO_2_ at 37^°^C. The transfection mix was then replaced with 200 μl complete medium before placing the plates back to the RTCA system. Cells were then incubated monitoring the cell impedance index every 15 min for 200 sweeps in first stage, 30 min for 200 sweeps in second stage, and continued at 60 min intervals for 100 sweeps in final stage. Time series cell impedance indices were extracted at regular time intervals. Negative controls were tested across each plate used to screen known (39) and potentially novel cell cycle-associated genes (22). At the time of screen a number of the known genes were considered uncharacterised. Each assay was based on results gained from running three replicate assays per plate, and repeated on three separate runs. The raw dataset was exported as a cell impedance (CI) index with rows named by N time points (measurement time points at 30 min intervals following the transfection) and columns named by well of samples. Statistical analysis of the data was performed using the R package “RTCA” to transform cell-impedance values into cell-index growth rate (CIGR) at regular time intervals during the measurement time ^44^. For scoring of the effect of gene silencing-induced proliferation arrest, the package “cellHTS2” was used to normalize average CIGR across samples^45^.

The library of esiRNAs (endoribonuclease-prepared short interfering RNAs, Sigma-Aldrich) employed here has been described elsewhere^46,47^. Negative control esiRNA reagents against sucrose isomaltase (*SI*), a gene not expressed by fibroblasts and collagen 1A2 (*COL1A2*), a gene expressed by fibroblasts but not associated with cell division. All control esiRNA reagents were used in each assay to verify the lack of non-specific effects of esiRNA treatment.

### Clone preparation

Entry clones in pDONR223 and containing open reading frames for candidate genes were sourced from the ORFeome collection ^48^. 50-150 ng of each entry clone was combined with 150 ng destination vector pcDNA-DEST47 or pcDNA-DEST53 (Life technologies) and 2 μl LR Clonase II enzyme mix (Life technologies). The reaction was incubated at 25^°^ C for 1 h. 1 μl (2 μg/μl) proteinase K was added to terminate the reaction, incubating for 10 min at 37^°^ C. 2 μl of the recombination reaction was added to chemically competent DH5α bacterial cells on ice and incubated for 20 min. DH5α cells were subjected to heat shock for 45 sec at 42^°^ C followed by 2 min on ice. 1 ml SOC medium was added and incubated at 37^°^ C with aeration. Cells were centrifuged at 2,000 rpm and resuspended in 100 μl LB before plating out on LB plates with 100 μg/ml ampicillin. Plates were incubated at 37^°^C overnight.

### DNA transfection and Confocal Microscopy

Transfection of HEK293 cells with Gateway destination clones and K2 transfection system (Biontex Laboratories, Munchen, Germany) was performed following manufacturer’s instructions. Following optimisation studies, 1 μg of expression plasmid and 2 μl of the transfection reagent were diluted in 500 μl in each well (24-well plate). For GFP fluorescence protein imaging, cells were fixed in 4% paraformaldehyde and labelled for 30 min with Texas RedX Phalloidin (1:40) (Invitrogen) and then stained for 5 min in 300 nM DAPI. For centrosomal staining, polyclonal anti γ-tubulin antibody (Sigma-Aldrich) was used. Alternatively, a polyclonal antibody anti α-tubulin (Abcam, Cambridge, England) was used to stain microtubules during the formation of mitotic spindles. Fixation was carried out by applying 300 μl of cooled methanol per well for 2 min on ice. Cells were washed three times with PBS and then blocked with 5% goat serum (Sigma-Aldrich) and 0.1% Triton in PBS for 1 h. Primary antibodies were then diluted accordingly in blocking solution, applied on coverslips and incubated either overnight at 4°C or for 2 h at room temperature. Cells were then washed three times for 1 h with PBS. The secondary antibody was diluted in blocking solution (1:500), applied on coverslips and incubated for 1-2 h at room temperature. Alexa Fluor^®^ 594 raised in donkey anti-rabbit IgG (H+L) (Life Technologies) was used as secondary for both primary antibodies, since they were used separately. Fluorescence images were captured on a Nikon EC-1 confocal microscope using Nikon EZ-C1 software. The following laser/filter combinations were used: DAPI nuclear stain (excitation 405 nm, emission BandPass 460/50 nm), eGFP (excitation 488 nm, emission BandPass 509 nm) and Texas Red X Phalloidin (Invitrogen) (excitation 543 nm, emission BandPass 605/70 nm).

## Results

### Identification of cell cycle-regulated genes in primary human fibroblasts

Two time-course microarray experiments were performed on populations of normal human dermal fibroblasts (NHDF) synchronised by serum starvation, as used previously for such studies^13,49^. Partial cell synchronisation was confirmed by flow cytometric analysis of propidium iodide-stained cells. Following serum starvation approximately 40% fewer cells were in the DNA replication phase (S) than in unsynchronised populations, and 24 h after the re-addition of serum the number of cells undergoing division had increased by 3-4 fold (S and G2/M phases) relative to the starved state (Fig 1A). Data derived from two transcriptomics experiments, one monitoring the cells every 6 h for 48 h following release from starvation, the second every 2 h over a period of 24 h, were subjected to quality control and corrected for batch variation. The datasets were combined and analysed together using Graphia^Pro^, a tool designed to analyse numerical data matrices into correlation networks^50^. A sample-to-sample correlation network confirming the correspondence between the two experiments and time-dependent transcriptional changes is shown in Fig 1B.

**Fig 1.**
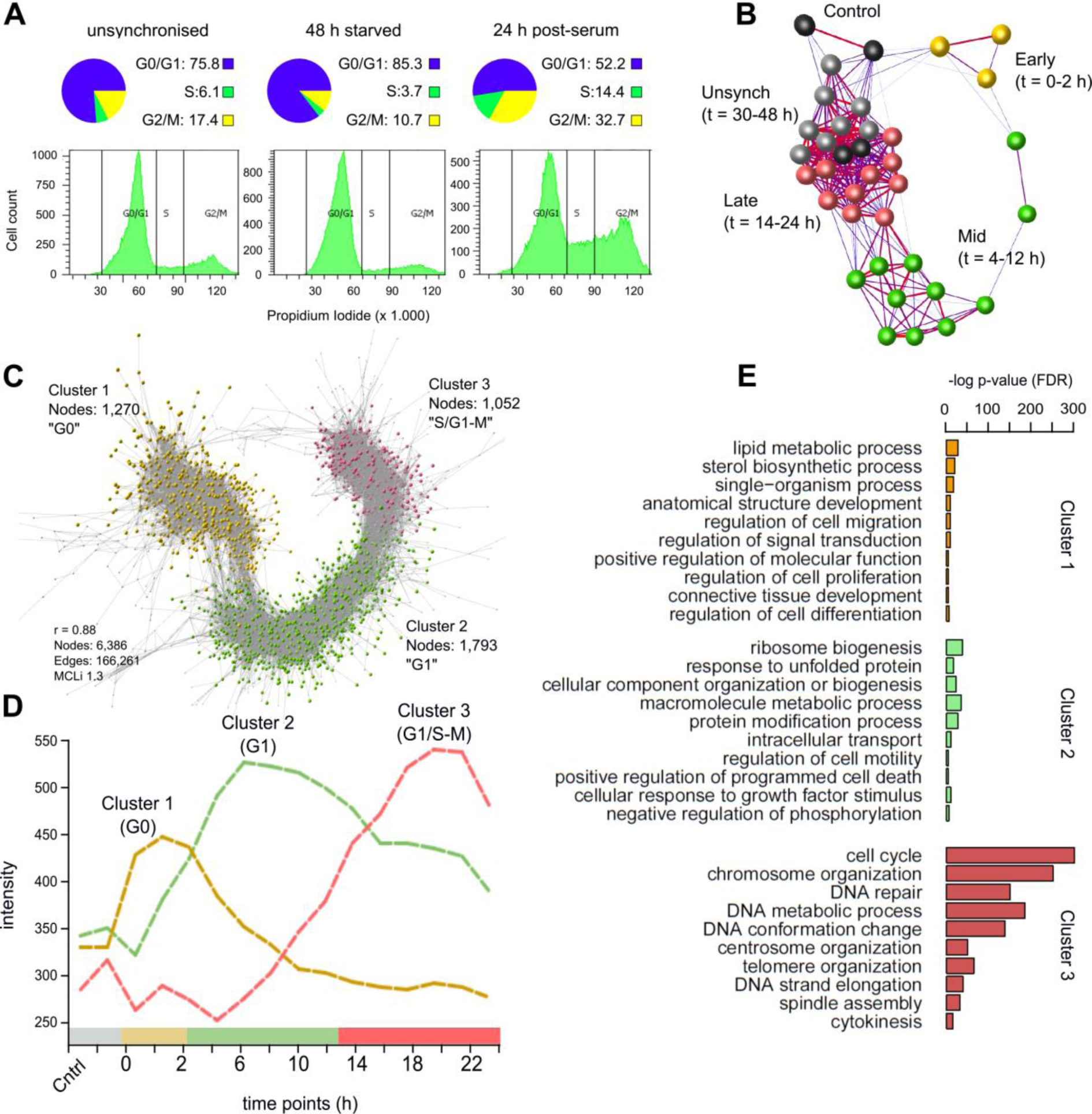
Network analysis of synchronised human fibroblasts (NHDF). **(A)** Flow cytometry data monitoring fibroblasts entering proliferation after serum refeeding. In control samples 76% of cells are in G_0_/G_1_ (“unsynchronised”) but following 48 h of serum starvation the figure had increased to 85%, whilst the proportion of cells in S/G2-M is decreased. 24 h post-serum 47% of cells were traversing S/G2-M phase (over 3 times greater than starved populations). **(B)** Sample-to-sample correlation graph where nodes represent individual samples. Samples of starved cells (0 h) and early proliferative populations (1-12 h), form distinct sub-groupings in the network, with a clear progression from early to late time points. Synchrony is lost at later time points, and samples group with unsynchronised populations. **(C)** Correlation graph of the transcriptional network of synchronised fibroblasts from a quiescence through to mitosis. The graph divides in three large clusters: NHDF_C1 (yellow) corresponds to genes whose expression is greatest in quiescent fibroblasts and decreases during the entry into mitosis (G_0_); NHDF_C2 genes (green) expression is associated with G1, their expression peaking between 1 and 8 h after the addition of serum; and the expression of genes in NHDF_C3 (red) start to rise from the beginning of the G1/S transition to mitosis. Nodes represent individual probesets. **(D)** Corresponding average expression profiles of genes in NHDF_C1-3. **(E)** GO enrichment analysis on the gene content of the three clusters.

A gene-to-gene correlation network was then generated using a threshold of *r* ≥0.88. This value is well above the distribution of random correlations (S1A Fig) and set to minimise the number of edges, while retaining a large number of nodes (S1B Fig). MCL clustering^33^ of the network was used to define the main phases of transcription associated with the cell cycle, generating 23 gene clusters. The three largest clusters accounted for 96% of the genes (probesets): NHDF_C1 (G0; 1,270 nodes, 1,176 unique Entrez IDs), NHDF_C2 (G1; 1,793 nodes, 1,739 unique IDs) and NHDF_C3 (S/G2-M; 1,052 nodes, 963 unique IDs) (Fig 1C). The average gene expression profile of the three clusters over the first 24 h following the re-addition of serum is shown in Fig 1D. NHDF_C1 comprised of genes induced during the starvation period (0 h), but down-regulated soon after the re-addition of serum to the growth medium. The average expression of genes within NHDF_C2 peaked around 6 h post-refeeding, consistent with gap (growth) phase (G1)^51^. NHDF_C3 included genes which were induced between 12 and 20 h post re-feeding, many of which remained elevated in their expression at later time points. Enrichment analysis performed on each gene cluster reported highly significant GO_BP term enrichments for all three clusters, the most significant of which are shown in Fig 1E. NHDF_C1 (G0-associated) was highly enriched with genes involved in lipid metabolism, such as *lipid metabolic process* and *sterol biosynthetic process*, supporting the evidence that these pathways are activated to adjust cellular metabolism during the starvation period^52^ (Fig 1E). NHDF_C2 was enriched in biological processes associated with cell growth, such as *ribosome biogenesis, macromolecule metabolic process, cellular component organisation* or *biogenesis and intracellular transport* and included many of the known regulators of G1 including *E2F3* and *CDK6 ^53,54^*. Finally, NHDF_C3 was highly enriched with terms such as *chromosome organisation, DNA repair, centrosome organisation, telomere organisation, DNA strand elongation, spindle assembly* and *cytokinesis* (Fig 1E). Transcripts within this cluster plus all nodes immediately adjacent to them (n+1), representing 1,207 unique Entrez IDs, were then selected for further analysis. A more granular cluster analysis of this coexpression network (MCLi 2.2) is presented in S1C Fig and S2 Table.

### Refinement of the core cell cycle gene signature

To refine the candidate list of NHDF S/G2-M phase-associated genes and eliminate genes that may be specific to differentiated fibroblast function, we examined their expression using the FANTOM5 consortium promoter level CAGE data (HCF5), derived from more than 100 different primary human cell types (495 samples)^27^. The 1,207 genes identified in the NHDF data were mapped to the FANTOM5 dataset returning 3,145 promoters with expression >5 TPM (tags per million) in at least one sample. These data were subjected to network analysis (*r*≥0.5). The graph contained 2,889 promoters (nodes) and 175,516 edges from which the MCL cluster algorithm (MCLi = 1.7) also generated 23 clusters (Fig 2A). HCF5_C1 (1,230 promoters of 745 genes) was enriched for cell cycle-associated genes (Fig 2B). Of the others, only HCF5_C2 exhibited any enrichment for the GO_BP term ‘cell cycle’ but at a much lower significance (Fig 2B). The average expression of HCF5_C1 gene promoters was greatest in highly proliferative cell populations such as embryonic stem cells, epithelial cells and a population of CD34^+^ hematopoietic stem cells. In contrast, monocytes displayed minimal expression of these genes, reflecting the low rate of proliferation in these populations^55^ (Fig 2C, top profile). The remaining HCF5 clusters contained promoters with a diverse range of expression profiles (Fig 2C). Many genes within HCF5_C1 also included alternative promoters (254 genes, 526 promoters) with distinct expression profiles that clustered independently (Fig 2D). Highlighted are six genes with known functions in the cell cycle, three of which, *RFC2, MCM5* and *MCM7* encode proteins known to be required for DNA replication^56^. The alternative promoters were most highly-expressed in immune cell types (Fig 2E).

**Fig 2.**
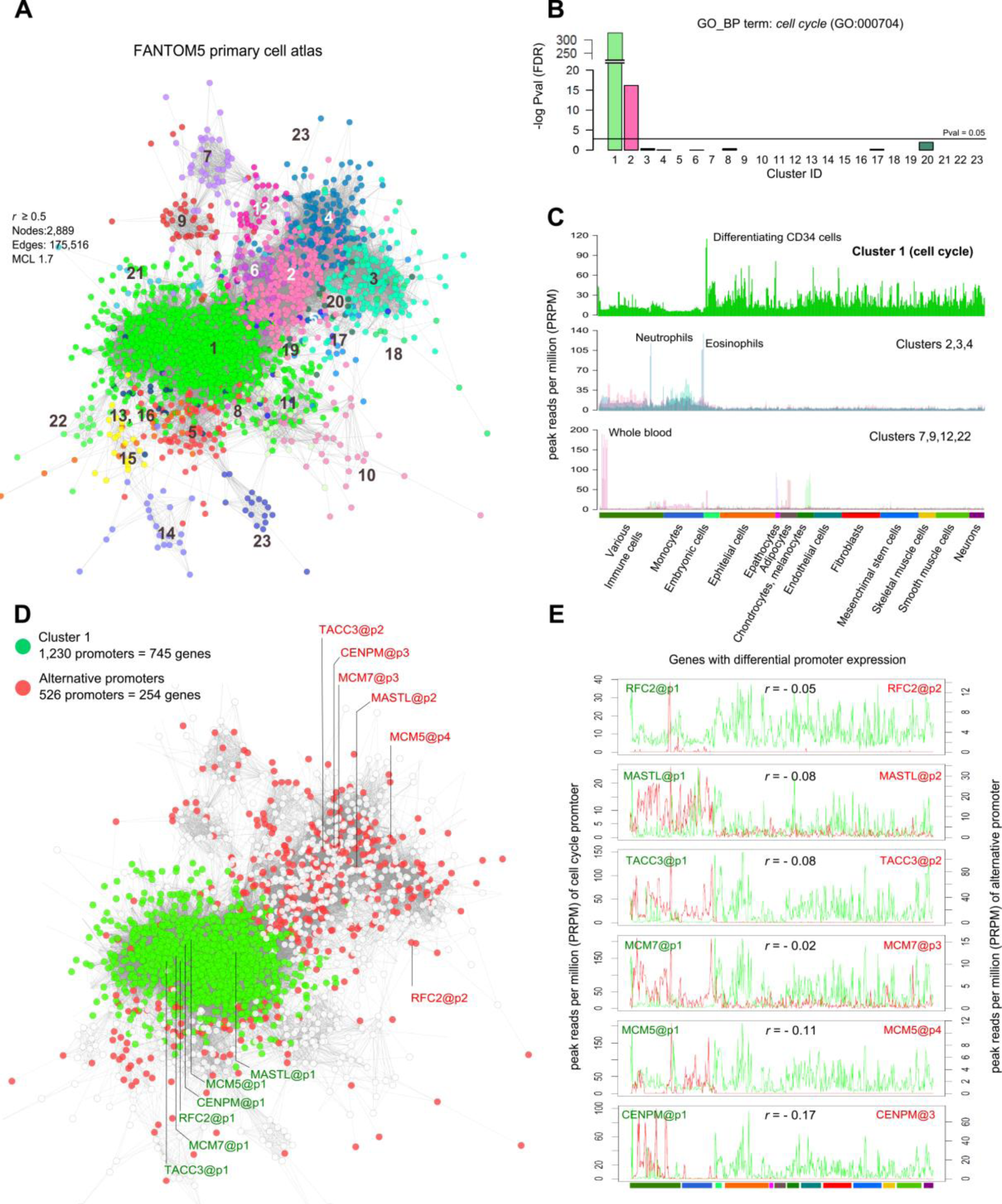
Co-expression of promoters associated with S/G2-M fibroblast genes across the FANTOM5 primary cell atlas. **(A)** Clustered graph representing the promoters of the S/G2-M phase associated genes identified in the NHDF data and their correlated expression in the context of the FANTOM5 primary cell atlas. Nodes represent individual promoters, their colour representing membership to co-expression clusters. **(B)** GO enrichment analysis for the GO_BP term *cell cycle* on each of the 23 clusters identified, cluster 1 to be highly enriched in cell cycle genes. **(C)** The expression profile of the HCF5_C1 promoters showed them to be transcribed in a wide variety of primary cells with highest expression in embryonic cells and a number of epithelial cells, but a relatively low expression in monocytes (top). In contrast other clusters, not enriched in cell cycle gene promoters, exhibited a different pattern of expression. The average expression of clusters HCF5_C2, 3 and 4 promoters was greatest in immune cell populations (middle). Others (bottom) exhibited cell type-specific expression, e.g. hepatocytes (HCF5_C7), adipocytes (HCF5_C9), whole blood (HCF5_C12) and melanocytes (HCF5_C22). **(D)** Nodes in the graph shown in A, were color-coded to show differential promoter expression. HCF5_C1 (green nodes) is comprised of 1,230 promoters corresponding to 745 genes, the red nodes represent an additional 526 promoters associated with 254 of the HCF5_C1 genes. **(E)** Promoter expression profiles of six genes being driven by promoters associated with the cell cycle (green profile) and expression of their alternative promoters (red profile).

Based upon the merge of the two datasets, the initial list of 963 genes (1052 probesets) generated from the analysis of NHDF cells, was reduced to a list of 745 genes with promoters in HCF5_C1 from the FANTOM5 data where the expression was tightly correlated across diverse human cell populations.

### Manual curation of the S/G2-M cell cycle list

The 745 genes identified above were individually curated. We removed 198 genes that were induced late in G1 and in advance of the likely onset of S phase. Conversely, we restored 132 genes, where the literature or other data (see below) indicated that they function in the cell cycle and individual examination of the FANTOM5 data indicated that they were, indeed, relatively more highly-expressed in proliferating cells, albeit not included in HCF5_C1.

To examine the inter-species conservation of the S/G-M transcriptional network, an additional transcriptomics experiment was performed on synchronised mouse embryonic fibroblasts (MEF). The majority of mouse/human orthologues showed a conserved expression pattern across the cell cycle (Fig 3A). The transcriptional network of the mouse fibroblast data was similar in topology to the NHDF data (S2A-B Fig) and an additional 22 known cell cycle genes were observed to co-cluster with the S/G2-M phase genes in these cells.

**Fig 3.**
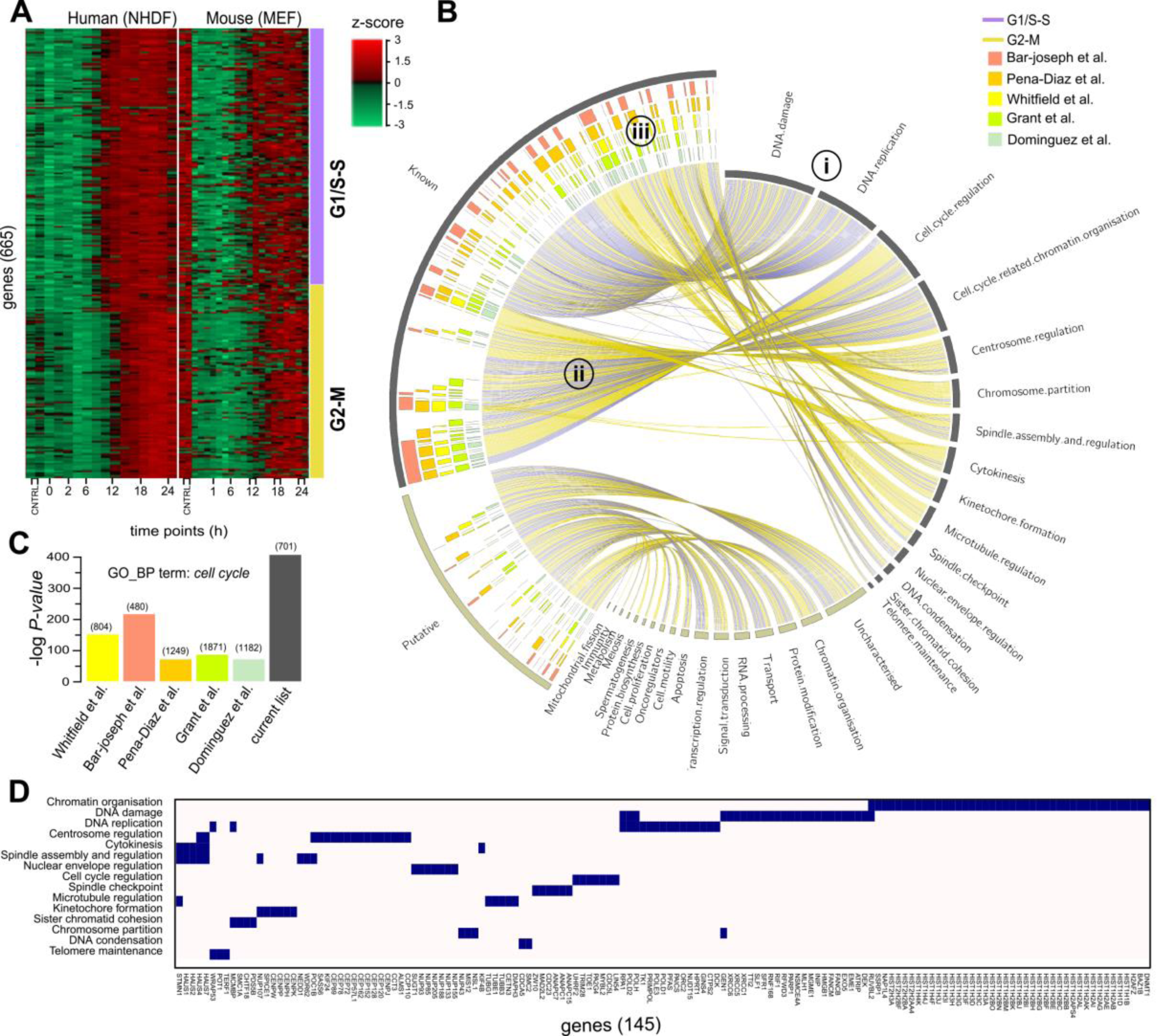
Analysis of the S/G2-M transcriptional network. **(A)** Heatmaps demonstrate a highly conserved pattern of expression between human S/G2-M phase associated genes and their 667 mouse orthologues over the first 24 h in human fibroblasts (NHDF) and mouse embryonic fibroblasts (MEF) following serum refeeding. Genes were ordered by the phase assignation calculated from the NHDF data. **(B)** CIRCOS plot showing the associations between the 701 S/G2-M human genes identified here and **(i)** the functional category with which they have been manually curated to belong. They have been divided as the whether they are ‘known’ or ‘putative’ cell cycle genes. **(ii)** Edges are coloured based on the phase assigned from the NHDF data. **(iii)** The inner coloured blocks show genes reported by previous human cell cycle transcriptomics studies. **(C)** Histogram of GO enrichment scores for the GO_BP term *cell cycle* for the current and previously published cell cycle lists. **(D)** Block diagram showing the functional category assignment of the 145 genes reported in the literature to be cell cycle-associated, but undetected by previous transcriptomics cell cycle studies.

The merged outcomes of the comparative analysis and manual curation of these data produced a set of 701 cell cycle regulated genes (see S2 Fig and S2 Table for a detailed cluster analysis of these data). The genes were then assigned to either ‘S’ or ‘G2-M’ phases by correlating them with the expression of known cell cycle phase-specific factors: *CDC25A* and *BRCA1* (S phase), and *CDK1* and *CCNB1* (G2-M phase)^57-59^. Accordingly, 380 genes were assigned to S phase and 321 to G2-M phase (S3A Fig). The two sets of phase-associated genes were analysed for enrichment of known binding motifs. Both sets were significantly enriched with cell cycle transcription factor binding sites. Amongst others S phase genes were shown to be highly enriched for E2F sites (10^−59^) and the G2-M genes for CHR (10^−15^) and NFY (10^−24^) sites (for detailed results see S3B Fig). Phase annotation was also consistent overall with those of previous cell cycle studies (S3C Fig).

After a systematic database and literature-based curation of the gene list, the majority (496) were found to be functionally associated with a cell cycle-related process (Fig 3Bi). For example, *DNA damage* and *DNA replication* linked predominantly with S phase annotated genes, and *Chromosome partition* and *Spindle assembly and regulation* being associated mainly with G2-M phase (Fig 3Bii). Other categories included a similar number of genes induced at either phase, such as *Cell cycle regulation.* For 205 genes little or no direct evidence of a direct involvement in the cell cycle could be found, although in some cases there was circumstantial evidence to support this relationship, e.g. publications showing their expression to be elevated in cancer. These genes are classified as ‘putative’ cell cycle genes. Others in this category encode proteins that function within pathways that potentially relate to cell division, e.g. apoptosis, whilst the association of yet others would appear more tenuous, e.g. RNA processing and immunity. For 53 genes, no functional information was found.

There is a significant overlap between our S/G2-M gene list and the cell cycle gene lists generated by previous studies (Fig 3Biii). However, the current list showed a greater enrichment of genes with the GO_BP term *cell cycle* when compared with published cell cycle lists (Fig 3C). Indeed, many well-validated S/G2-M phase genes (145) were not shown to be regulated in any of the previous transcriptomics study, including mitotic regulators such as *MADL2L2,* four members of the augmin complex *HAUS1,2,4,7*, three members of the APC/C cyclosome complex, *ANAPC1,7,15,* multiple DNA replication-dependent histone isoforms (36) and several genes encoding components of the centrosome (*CEPs)* and the nucleopore complex (*NUPs*) (Fig 3D). Conversely, there were 345 genes annotated with the GO_BP term *cell cycle* identified by at least one of the five previous human transcriptomics studies but not the current study. When examined in the context of the current NHDF data, many were induced during G1, whereas others did not show significant variation in their expression over the cell cycle (S4 Fig). As noted above, we have deliberately excluded genes that are known to be induced in G1, although this gene set may include genes that are essential for cell cycle progression^26^.

A table of all 701 genes, which includes eight pseudogenes and three non-coding RNAs, with corresponding classifications and evidence supporting their functional association with cell cycle can be found in S1 Table. A simple confidence score was calculated for all genes in the list based on available experimental evidence from this and previous studies.

### Validation of the S/G2M transcriptional network using independent data

To further validate the conservation of co-expression of the S/G2-M gene list, we explored two additional datasets. The first was a human tissue expression atlas (HTA) of RNA-seq data derived from 95 samples of 27 human tissues^32^. Of the 701 S/G2-M genes defined here, 655 were identified in these data and their co-expression examined. At a correlation of *r* ≥ 0.5, 641 genes were present in the graph, which divided into two MCL-defined clusters encompassing 549 genes (Fig 4A). The genes in these clusters were expressed widely, with an elevated expression level associated with proliferative tissues (Fig 4B). HTA_C1 was comprised of genes whose expression in the testis was higher (Fig 4B, top) as compared to the expression of HTA_C2 genes, which showed highest expression in bone marrow and lymph node (Fig 4B, bottom). The majority of known S/G2-M genes clustered together and significantly, so did the putative cell cycle genes (Fig 4C-D), supporting their association with this system. A second promoter-level dataset produced by the FANTOM consortium (MDF5)^27^, comprised of 17 mouse tissues sampled at multiple intervals during embryogenesis and post-neonatal development. Again the data for only the S/G2M genes (2,141 promoters mapping to 658 genes) was examined. Similarly, the majority of the promoters/genes co-clustered, with the exception of a few small clusters (Fig 4D). In general the promoters in MDF5_C1 exhibited highest expression in embryonic tissues, their expression markedly decreasing with developmental age, a pattern reflecting the reducing rate of proliferation during development (Fig 4F, top profile). A notable exception to this was in the case of the spleen and thymus, where expression peaked around birth. In line with observations in the human tissue atlas dataset, a portion of S/G2-M genes (MDF5_C2) were predominately expressed in adult testis (Fig 4F, middle profile). Multiple smaller clusters, the majority of which were associated with alternative promoters of cell cycle-associated genes, exhibited tissue-specific promoter expression (Fig 4F, bottom profile). Again putative S/G2M genes were co-expressed with the known cell cycle genes (Fig 4G-H). Co-expression of the S/G2-M genes within the context of all genes are shown in S5A-B Fig, for the HTA dataset and in S5C-D Fig for the MDF5 dataset. Promoter analysis of the 205 putative cell cycle genes alone showed that they were enriched in known S/G2-M transcription factor binding sites, i.e. for E2Fs and NFY, further supporting their associated with cell division (S5E Fig).

**Fig 4.**
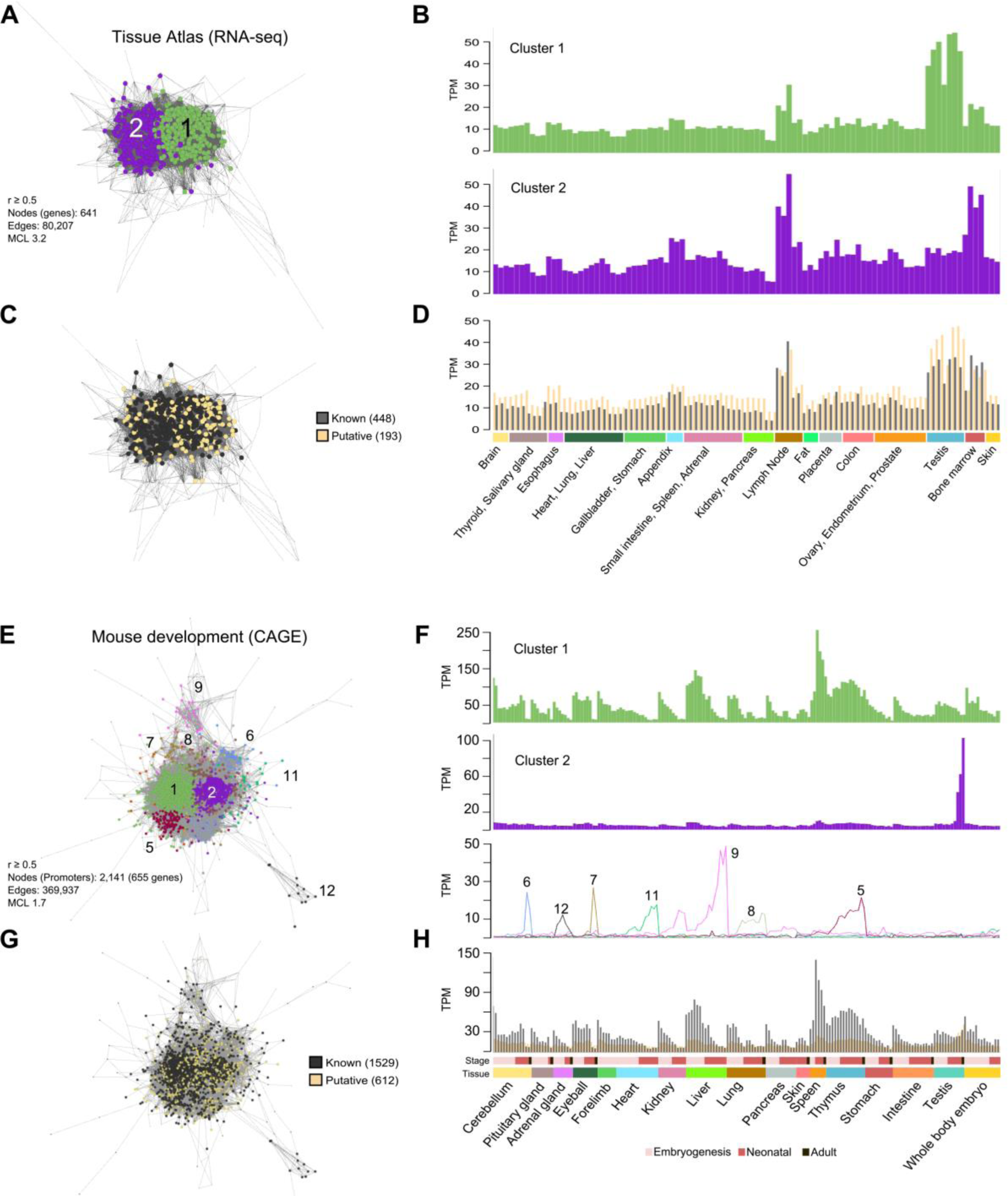
Confirmation of coexpression of S/G2-M genes across human and mouse tissues. **(A)** Clustered coexpression network of S/G2-M genes across human tissue atlas (HTA). **(B)** The average expression profile of the genes in the two main clusters is very similar with the exception that genes in HTA_C1 are strongly expressed in the testis. **(C)** Interesting both known and putative cell cycle genes cluster together, **(D)** having very similar expression profiles. **(D)** Clustered coexpression network of promoter level data of mouse orthologues of human S/G2-M genes in the mouse development dataset from FANTOM5 (MDF5). Here a number of clusters are observed. **(F)** The largest group (MDF5_C1) are highly expressed in all developing tissues but expression levels generally decrease with age. However, in the case of spleen and thymus highest levels of express are observed around birth. MDF5_C2 promoters are highly expressed in adult testis and alternative promoters that form the majority of other clusters show a variety of tissue-specific expression patterns. **(G-H)** Promoters for known and putative cell cycle genes cluster together and exhibit a similar expression profile.

### Experimental corroboration of the uncharacterised S/G2-M genes

Many gene products required for S/G2-M phase, are localised to specialised cell cycle-associated organelles or structures. For example, chromosome segregation during mitosis requires formation of kinetochores at centromeres and the correct attachment of kinetochores to spindle microtubules emanating from microtubule organising centres, e.g. centrioles and centrosomes. Accordingly, we tested the subcellular localisation of 28 candidate genes by cDNA transfection in HEK293 cells. As positive controls we included CENPA, TACC3, DONSON and MGME1^60-63^, the localisation of which were confirmed by these assays (S1 Data). Each ORF was tagged with GFP at both the C-and N-terminals^48,64^. After inspection of the expression of the 56 protein constructs (two per clone), their subcellular localisation was analysed (S2 Data). As summarised in Fig 5A, nuclear localisation was the most frequently observed (15/28) followed by centrosomal-like localisation (11/28) and cytosol (9/28). In some instances localisations were congruent with organelles such as the ER, Golgi apparatus, vesicles and mitochondria, possibly representing non-specific protein deposits. In around half of cases, the C- and N-terminal tagged proteins produced the same localisation (Fig 5A). No cases of completely discrepant localisations between the two constructs were observed. For the 11 constructs showing centrosomal-like staining, we examined their expression along with centrosomal marker γ-tubulin. Of these, C18orf54, C3orf14 and CCDC150 clearly co-localised with γ-tubulin during different mitotic stages, i.e. prophase, prometaphase and metaphase (Fig 5B-D). For the other constructs co-localisation with γ-tubulin was not demonstrated (not shown). Confocal images of all 28 proteins screened can be found in S1 Data.

**Fig 5.**
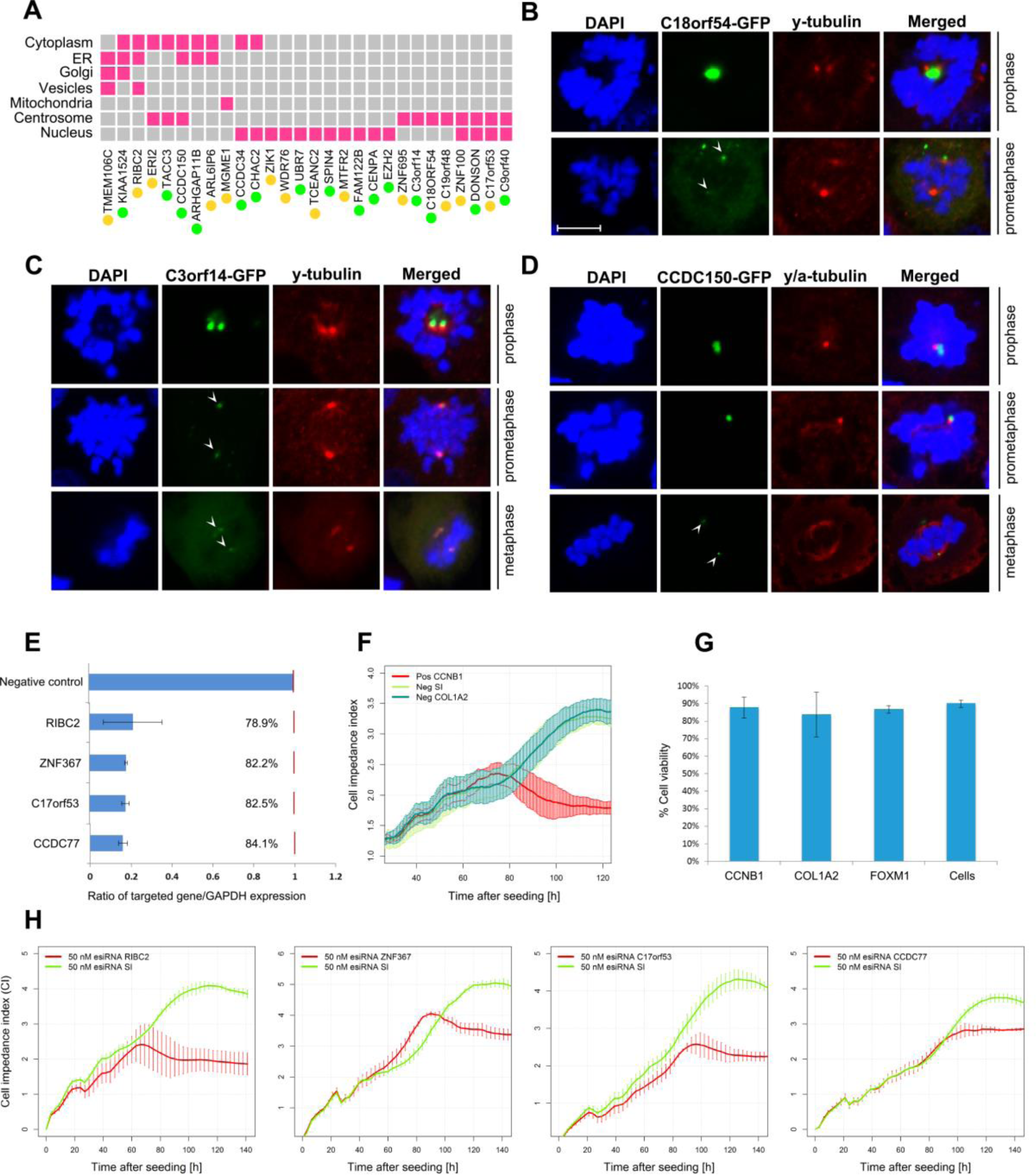
Subcellular localisation and RNAi assays of candidate cell cycle components. **(A)** Matrix summarising the subcellular localisations of the 28 proteins screened. For a full description of these data see S1 Data. **(B-D)** C3orf14, CCDC150 and C18orf54 were found localised on the centrosomes at different stages of mitosis. Proteins are tagged with GFP (green), nuclei are stained with DAPI (blue), and centrosomes marked with anti γ-tubulin antibody (red). Scale bar = 10 μm. **(E)** Knock-down efficiencies of siRNA against four potential novel cell cycle genes measured as the ratio between the silenced gene expression and GAPDH expression. **(F)** Positive control cyclin B (*CCNB1*) silencing decreased cell impedance index (proliferation) compared to negative controls for sucrase-isomaltase (*SI*) and collagen 1 A2 (*COL1A2*). **(G)** Viability assays after gene knock-down of two known cell cycle regulators (*CCNB1* and *FOXM1*) and a negative control (*COL1A2*) compared to untransfected cells. **(H)** Proliferation profiles following gene knock-down of four uncharacterised but putative cell cycle genes *RIBC2, ZNF367, C17orf53* and *CCDC77* compared to knock down of *SI*. For a full description of the results all knock-down experiments performed here, see S2 Data.

To examine whether reducing the expression of the novel S/G2-M phase genes affected cell proliferation, we tested the effect of mRNA knockdown in human fibroblasts using esiRNAs. We achieved around 80% knock-down efficiency in all cases examined (Fig 5E). As a positive control, knockdown of cyclin B1 (*CCNB1*) produced a strong inhibition of cell proliferation compared to control esiRNAs (Fig 5F) and transfection had no effect on cell viability (Fig 5G). Of the 39 knockdowns of known cell cycle-regulated genes tested, only twelve (*ARHGAP11A, CCNB1, CCNE1, CENPA, CEP85, ESPL1, FAM111A, FIGNL1, FOXM1, KIF11, MAD2L1, REEP4*) had a significant impact on the rate of cell proliferation. Similarly, of the 22 uncharacterised cell cycles genes 10 (*C17orf53, CCDC77, DEPDC1B, FAM72B, GSTCD, NEMP1, RIBC2, RPL39L, UBR7, ZNF367*) significantly inhibited proliferation (Fig 5E; results of these analyses in S2 Data). The Mitocheck database is a resource listing the cellular phenotypes from a genome-wide RNAi-screen of human proteins, recorded by high-throughput live cell imaging^8^. Of the known gene components listed for which results were available, 136/490 (27.8%) resulted in one or multiple cell phenotypes pointing to cell cycle defects. Amongst the candidate cell cycle genes, 12.8% were associated with a cell cycle phenotype. For example, *ZNF85* silencing led to abnormal chromosome segregation and mitotic metaphase plate congression, knock-down of *ZNF90, UBALD2* and *CCDC34* led to cell death and *ZNF738, CCDC150* and *ZNF788* knockdowns resulted in abnormalities in the size and shape of nuclei. Mitocheck results have been added to the gene list presented in S1 Table. Finally, ToppGene^65^ was used to search for phenotypes significantly associated with mutations in the S/G2-M genes. In man, 79 phenotypes were identified, the most significant of which included embryonic growth defects, e.g. microcephaly, growth retardation and various cancers. In mouse, 242 phenotypes were recorded as enriched, the most significant were abnormal cell cycle, embryonic lethality and abnormal nuclear morphology, again supporting a strong association with cell cycle defects. A full list of the phenotypes enriched for this list and the genes associated with them is provided in S3 Table.

## Discussion

Mitotic cell division is perhaps the most fundamental of all biological processes and functional orthologues of many of the core components are conserved across species. Curated databases list and classify the function of cell cycle components^66^ and place them into pathways^39,67^. In every system studied, from yeast to man, there are numerous genes required for cell division that are transcriptionally regulated and associated with a given phase of the cell cycle^17,68^. It could be argued that all genes involved in anabolic processes are cell cycle-related, since an increase in cell size is usually precondition for cell division. Similarly, genes regulating entry into the cycle, e.g. growth factors, are often considered to be cell cycle proteins. In the context of this work, we use the term to refer only to the set of proteins that are required when a cell commits to undergo mitosis^20^. Accordingly, we have sought to define the core set of cell cycle genes expressed during mammalian S/G2-M, demonstrating them to form a highly correlated transcriptional network across tissues and cell types. As a gene signature, S/G2-M genes effectively define the mitotic index of a cell population.

The gene expression patterns observed here in fibroblasts were broadly consistent with previous studies using the same cell type and synchronisation method^13,49^. However, a wound-healing response, triggered by the serum, may confuse efforts to identify cell cycle-related transcripts in fibroblasts^18^. To circumvent this issue, we complemented our analysis by examining the co-expression of the fibroblast-derived S/G2-M associated genes using the FANTOM5 primary cell atlas^27^ to remove genes that showed evidence of cell-specificity in their expression. These analyses were further refined by comparison to expression studies in synchronised mouse fibroblasts and detailed examination of the primary data. The result is a list of 701 S/G2-M-regulated genes, which are highly enriched in relevant GO terms and transcription factor binding sites. Based on manual curation of published reports, 496 of these genes encode ‘known’ cell cycle proteins, many listed as such in annotation databases e.g. GO and UniProtKB. This list partially overlaps with the findings of previous transcriptomics studies on human cells but interestingly, transcriptional regulation of 145 of the known S/G2-M associated genes was not detected in any of the earlier reports^13-16,18^. The majority of previous studies sought to define cell cycle genes as having a wave-like expression profile over multiple rounds of cell division, using Fourier transform-based methods to identify them. However, cell division in populations of cells rapidly becomes asynchronous, and a fraction of them do not commit to a second cycle ^13^. In the current study, this fact was reflected in the loss of synchrony in the cell cycle gene expression signature after 30 hours, consistent with FACS analyses (data not shown). Not only did previous studies exclude many bona fide cell cycle genes the different criteria and analytical methods used produced a poor consensus^15^. The correlation-based network approach used in this study is a more efficient way to identify phase-specific cell cycle genes^20^.

The strong association of the many putative cell cycle genes identified here was further demonstrated by their conserved coexpression across adult human and developing mouse tissues. Some of the S/G2-M phase genes we identified have only been validated relatively recently. For example, *PRR11* (proline rich 11), mutations in which have been associated with cancer, was shown to regulate S to G2-M phase transition^69^. Links to cancer biology also suggests function for two of the three lncRNAs identified by this study, *DLEU1*&*2*^70,71^. There are nine genes annotated as being involved in apoptosis, a process that can be initiated if a cell fails a mitotic check point. *CASP2,* long considered to be an orphan caspase^72^, is recognised as a key factor in driving cell apoptosis (mitotic catastrophe) triggered by mitotic abnormalities, such as defects in chromosomes, mitotic spindles, or the cytokinesis apparatus^73,74^. Other genes within the list await functional validation.

Amongst the 205 putative cell cycle genes, there are 53 complete functional orphans. Fourteen of the 27 we tested localised wholly or partially to the nucleus and 11 showed evidence of centrosomal localisation, an organelle vital for cell cycle progression^75^. Another three, CCDC150, C3of14 and C18orf54 co-localised with γ-tubulin (a centrosomal marker). A recent study confirmed this localisation for C3orf14^76^. The sub-cellular localisations reported here were in many cases also supported by IHC results reported by the Human Protein Atlas database^37,37^ (data not shown). RNAi knockdown assays were also performed on a range of known and uncharacterised genes from the list. In these assays, 10 of the 22 uncharacterised proteins showed differences in the rate of cell proliferation following gene knockdown, suggesting non-redundant functions in cell proliferation, with a hit rate slightly higher than the known cell cycle genes tested (Fig 5H). Taken together, these validation data suggest that the large majority of the novel cell-cycle regulated genes identified here will be found to function in some aspect of S/G2-M biology.

The FANTOM5 human and mouse promoterome data provide definitive locations for the transcription start sites of genes. Of the 701 genes identified here, in the primary cell atlas data at least 254 use alternative promoters that drive their expression outside of the context of the cell cycle. Among them, three are involved in the assembly of the replisome, namely: *MCM5, MCM7* and *RFC2*^56^, and had significant expression from alternative promoters in certain immune-related cell populations. This observation is in accordance with a previous report showing that factors of the minichromosome maintenance complex (MCMs), including MCM5 and MCM7, were found to be present on the IRF1 promoter in STAT1-mediated transcriptional activation, when cells were treated with IFN-y^77^. MCM5, in particular, was shown to directly interact with STAT1 and to be necessary for transcriptional activation^78^. Similar observations were made in the analysis of the mouse development time course data, where many bona fide cell cycle proteins are strongly expressed in the testis, where they may play a role in meiosis or be part of the centrosomal biology associated with flagella. The ‘moonlighting’ of cell cycle genes in other scenarios also likely breaks up the transcriptional signature in co-expression analyses across datasets comparing tissues or cells^21-25^. These alternative transcripts may be regulated in a unique manner to support DNA-dependent processes such as recombination, somatic hypermutation and class switching that are unique to leukocytes, or other distinct functions.

In summary, this study set out to define the transcriptional network associated with the final stages of the human cell cycle, between entry into S phase through to the completion of mitosis. The aim was not only to summarise the known components of this system but to identify new ones. Through detailed analyses of multiple human and mouse datasets we have defined 701 genes as being upregulated during the S/G2-M phase of the cell cycle, many of which are conserved across species. Based on promoter expression some proteins would appear to function exclusively within the context of cell division, others would appear to have additional roles outside of this system. Functional assays performed on a number of the uncharacterised genes strongly suggests that many are indeed novel components of the cell cycle machinery. The gene list provided represents the first comprehensive list of experimentally derived and validated S/G2-M phase associated genes. As such this work provides a valuable resource of both the known and potentially novel components that make up the many pathways and processes associated with mitotic cell division.

## Supplementary Captions

**S Fig1. Analysis of NHDF transcriptomics data. (A)** Plot shows the distribution of correlation values between 2,000 genes randomly selected from the NHDF data compared with that of the same genes but with permuted values. The threshold used for analysis excludes random correlations. **(B)** Plot showing number of edges and nodes as a function of the correlation coefficient. A threshold of *r* ≥ 0.88 was selected to include a minimal number of edges, while retaining a large number of nodes. **(C)** Network graph of the data clustered at MCLi 2.2 and average expression profiles of the main clusters showing gene expression as a function of time.

**S Fig2. Network analysis of the MEF data. (A)** Network graph of the MEF data (MCLi 2.2) at *r* ≥ 0.88. **(B)** Average expression profile of the major clusters of cell cycle reulated genes.

**S Fig3. Phase assignation analysis and comparison with previous data. (A)** The 701 genes associated with S/G2-M phase were assigned as being either ‘S’ or ‘G2-M’ phase according to their correlation with *bona fide* phase-specific cell cycle genes (see text), resulting in 380 S phase genes and 321 G2-M genes. **(B)** Motif enrichment analysis performed using HOMER were run on the two gene subsets returning significant enrichments for motifs bound by transcription factors known to be active in the corresponding phases. **(C)** Our phase assignation was compared to those of five previous studies. Heatmaps for each of these comparisons show overal consistent phase assignation.

**S Fig4. Expression of the 701 S/G2-M genes identified here and 345 other cell cycle-annotated genes.** Expression of the 701 genes show up-regulation associated with entry into S phase throgh to the completion of M phase. The majority of the other 345 genes identified by the five previous cell cycle studies^13,14,18,34,35^ but not in this study and associated with the GO_BP term cell cycle showed induction at earier time-points. Expression values were transfomed to z-score (see legend).

**S Fig5. Coexpression networks of human tissue (HTA) and mouse tissue development (MDF5). (A)** Clustered coexpression network of HTA analysis, showing clusters of genes exhibiting specific expression patterns. The location of the majority of cell cycle genes is highlighted by dotted red lines and in this graph reside in cluster HTA_C6. **(B)** Enrichment analysis shows this cluster to be highly enriched in genes with the GO_BP term cell cycle and with genes included in our list. **(C)** Clustered coexpression network of MDF5 analysis, showing clusters of genes exhibiting specific expression patterns. Cell cycle genes clusters are highlighted by dotted red lines. **(D)** Enrichment analysis shows clusters MDF5_C11, MDF5_C27 and MDF5_C1 to be significantly enriched with the GO_BP term *cell cycle* and with genes included in our list. **(E)** Motifs enrichments of ten randomly selected subsets of the known group of S/G2-M genes (left) equivalent to the number of putative S/G2-M genes (right). Significant cell cycle-associated TF, such as E2Fs and NFY, were identified in both cases.

**S Table 1. List of all known and putative S/G2-M phase associated genes identified by the current study.** The table contains an annotated list of 701 genes identified by this study and the evidence from the current and previous studies linking them to the cell cycle.

**S Table 2. Clustering results from network analysis of the NHDF and MEF cell cycle associated transcriptome.** Lists of genes whose expression is regulated at any point during the cell cycle in human and mouse of mouse fibroblasts.

**S Table 3. Phenotypes associated with mutations in S/G2-M phase genes.** ToppGene was used to analyse what human and mouse phenotypes are associated with the 701 genes identified here and statistically over-represented. This table provides a full list of these phenotypes and which genes they are associated with.

**S Data 1. Results from subcellular localisation studies of uncharacterised cell cycle proteins.** A summary of results from experiments using GFP-tagged proteins to study the localisation of a number of uncharacterised putative cell cycle proteins.

**S Data 2. Results of RNAi screens of uncharacterised cell cycle proteins.** A summary of the results from esiRNA knockdown studies of known and putative cell cycle genes.

## Acknowledgement

B.G. is a recipient of a BBSRC funded EastBio studentship. T.L. is supported by a Sir Henry Dale Fellowship jointly funded by the Wellcome Trust and the Royal Society (206211/Z/17/Z). The Wellcome Centre for Cell Biology is funded by Wellcome grant 203149/Z/16/Z. M.B., D.A.H. and T.C.F. are funded by an Institute Strategic Grant from the Biotechnology and Biological Sciences Research Council (BBSRC) (BB/JO1446X/1).

## Author contributions

B.G., M.B., T.R. and S-H.C. performed the majority of laboratory work described here. B.G. and T.C.F. performed the bioinformatics analyses and, B.G., T.L., S.W., D.A.H. and T.C.F. wrote and edited the manuscript. T.C.F. supervised the project and conceived of the idea behind the work.

